# Development of immortalized rhesus macaque kidney cells supporting infection with a panel of viruses

**DOI:** 10.1101/2022.07.18.500407

**Authors:** Stefanie Reiter, Sabine Gärtner, Stefan Pöhlmann, Michael Winkler

## Abstract

Non-human primate (NHP)-based model systems are important for biomedical research, due to the close phylogenetic relationship and physiologic similarities of NHP and humans. In infection research, NHP models are used to model various viral diseases including Ebola, influenza, AIDS and Zika. However, only a small number of NHP cell lines are available and generation of additional cell lines could help to refine these models.

We immortalized rhesus macaque kidney cells by lentiviral transduction with a vector encoding telomerase reverse transcriptase (TERT). Expression of kidney markers on these cells was analyzed by flow cytometry and quantitative real-time PCR (qRT-PCR) was employed to determine functionality of the interferon (IFN) system. Finally, we assessed susceptibility and permissiveness for virus infection by the use of pseudotyped particles and replication-competent viruses.

We report the generation of three TERT-immortalized cell lines derived from rhesus macaque kidney. The cell lines expressed the podocyte marker podoplanin and expressed MX1 upon stimulation with IFN or viral infection. Further, the cell lines were susceptible to entry driven by the glycoproteins of vesicular stomatitis virus, influenza A virus, Ebola virus, Nipah virus and Lassa virus. Finally, these cells supported growth of Zika virus (ZIKV) and the primate simplexviruses Cercopithecine alphaherpesvirus 2 (CeHV2) and Papiine alphaherpesvirus 2 (PaHV2).

We developed IFN-responsive rhesus macaque kidney cell lines that allowed entry driven by diverse viral glycoproteins and were permissive to infection with Zika virus and primate simplexviruses. These cell lines will be useful for efforts to analyze viral infections of the kidney in macaque models.

## Introduction

Due to their close phylogenetic relationship to humans and physiologic similarities, non-human primates (NHP) are important animal models for biomedical research. In infectious disease research, NHP models of human immunodeficiency virus (HIV), Ebola virus (EBOV) or Zika virus (ZIKV) infection mirror closely the pathogenesis in human patients (1–3). However, because of ethical and legal considerations, strategies for replacement, reduction and refinement (3R) of NHP models need to be implemented (4). As a consequence, alternative non-animal methods have been promoted, including but not limited to cell culture models.

Cell lines offer a simple system for analysis of viral infection that combines low cost and ability for upscaling (5). Usually cell lines are established by explantation of native or tumor tissue and selection of continuously growing cells. To overcome the limited reproductive capacity of native tissue cells, cells can be immortalized using viral or cellular genes like simian virus 40 (SV40) large T (6) or the catalytic subunit of telomerase reverse transcriptase (TERT) (7, 8).

In contrast to the broad availability of human cell lines only a handful of cell lines of NHP origin are available from commercial suppliers (such as ATCC), and only a few additional cell lines have been described in literature (9–11). Thus, there is a clear need for the development and characterization of new cell lines of NHP origin. We have established and immortalized, using TERT, novel cell lines derived from rhesus macaque kidney tissue. These cell lines expressed the podocyte marker podoplanin and expressed the IFN-stimulated gene (ISG) MX1 upon stimulation with IFN or viral infection. Finally, these cells were susceptible to infection by viruses from several families, as assessed by pseudotyped viral particles, and supported productive infection with primate herpesviruses and ZIKV.

## Materials and Methods

### Plasmids and oligonucleotides

Plasmids MLV-gag/pol, MLV-luc, Sgpdelta2 and pHIT/G have been described (12–14). Plasmid HIV-gag-pol was a kind gift from Alexander von Hahn. Plasmids for expression of glycoproteins from Vesicular stomatitis virus (VSV-G), Zaire ebolavirus (EBOV-GP), Nipah virus (NIV-F and NIV-G), influenza A virus strain WSN (IAV HA and NA) and Lassa mammarenavirus (LASV-GPC) have been described previously (14–18). Oligonucleotides (Table 1) were purchased from Sigma-Aldrich (Steinheim, Germany).

**Table 1:**
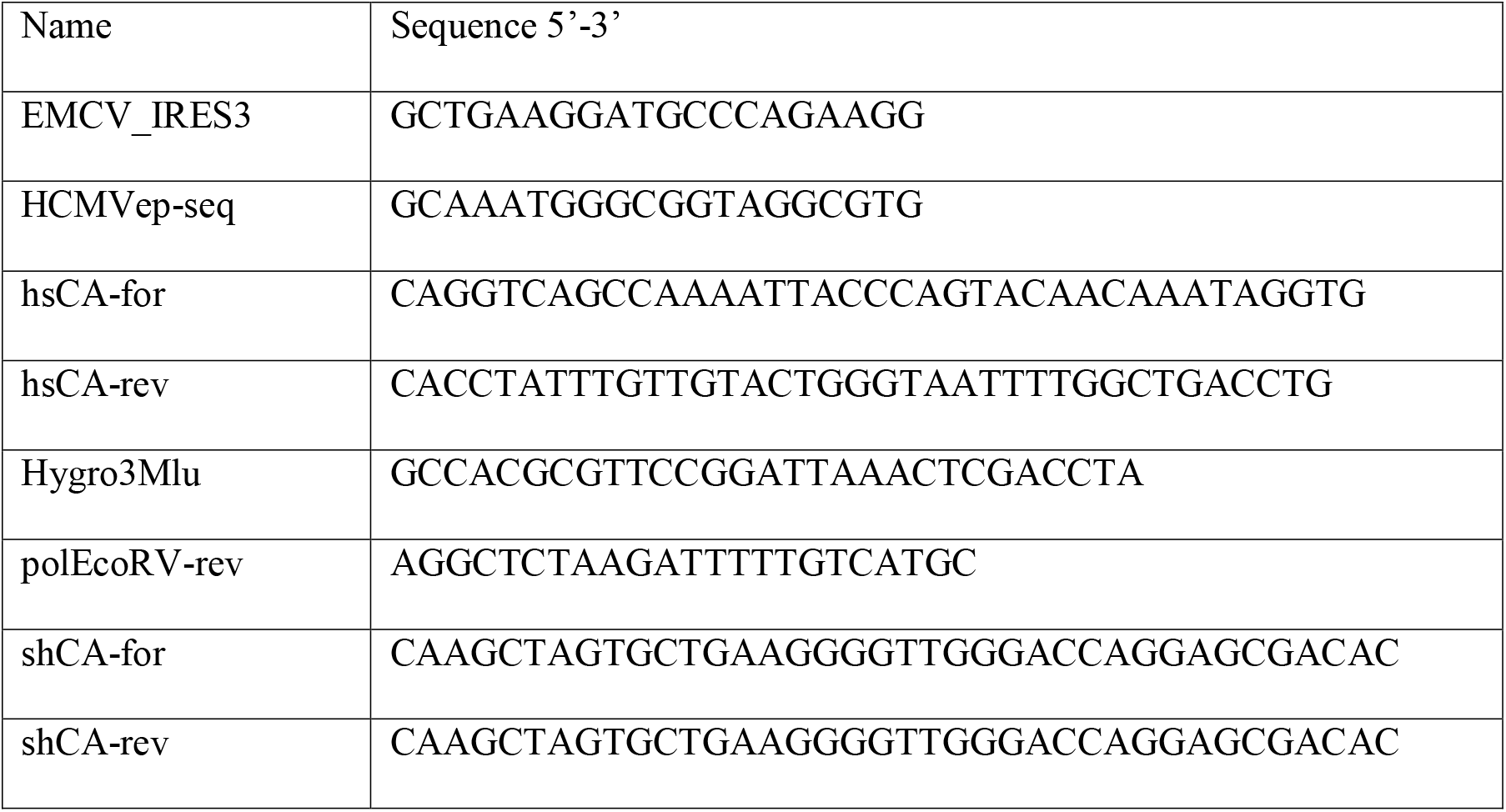
Oligonucleotides used for cloning.

Retroviral vector pQCXIHy-hTERT was generated by subcloning of hTERT as PmlI/SalI fragment from pBabehygro-TERT (kind gift from Parmjit Jat) (8) into pQCXIHy-mcs (19) cut with HpaI and XhoI. This vector has a shortened C-stretch at the 5’ end of the internal ribosomal entry site (IRES) and the modified multiple cloning site of pQCXIP-mcs (20) which was ultimately derived from pQCXIP (Clontech, Palo Alto, CA, USA). Finally, pQCXIP was modified by mutating an AarI site within the IRES sequence and including AarI, NheI and XbaI sites in the multiple cloning site, to give pQCXIdAP-mcs2.

For lentiviral transduction, a pReceiver-Lv205-based expression plasmid (Genecopoeia, Rockville, MD, USA) was modified stepwise to harbor a pQCXIP-based expression cassette. First, a NdeI/Pfl23II fragment from pQCXIP-PB1 was inserted into EX-A2639-Lv205 to give pLenti-PB1. Second, a NdeI/Acc65I from pQCXIdAP-mcs2 was inserted into pLenti-PB1 to replace the PB1 gene with a multiple cloning site. The resulting vector, termed pLenti-IP-mcs, contained human cytomegalovirus (HCMV) enhancer/promoter, multiple cloning site, internal ribosomal entry sequence and puromycin resistance gene. The multiple cloning site had restrictions sites for MunI-AgeI-PmlI-XbaI-XhoI-BamHI-NotI. To obtain pLenti-IHy, harboring a hygromycin resistance gene, the puromycin resistance gene was replaced by an Acc65I/MluI digested PCR fragment amplified from from pQCXIHy-mcs with primers EMCV_IRES3 and Hygro3Mlu. The human TERT gene was then subcloned from pQCXIHy-hTERT into pLenti-IHy-mcs as an AgeI/BspEI fragment to give pLenti-IHy-TERT.

To generate an HIV-based system with improved transduction of rhesus monkey cells amino acids 1-204 of the HIV capsid gene (hCA) was replaced with amino acids 1-202 of SIV capsid gene (sCA) to give HIV gag-pol(SCA) (21, 22). For this the flanking HIV parts were amplified by PCR using primer HCMVep-seq/hsCA-rev and shCA-for/polEcoRV-rev and HIV-gag-pol as template, while the sCA was amplified from Sgpdelta2 using primer hsCA-for/shCA-rev2. All the fragments were combined by splice overlap PCR and cloned as EcoRI/EcoRV into HIV-gag-pol.

### Cell culture

Cell lines 293T, Vero76 and A549 were cultivated in DMEM supplemented with 10% fetal calf serum (FCS) and penicillin/streptomycin. Identity of human cell lines was verified by short tandem repeat (STR) analysis (23). Hybridoma cell line D1-4G2-4-15 was purchased from LGC/ATCC (Teddington, UK) and cultivated in RPMI1640 supplemented with 10% FCS and Pen/Strep.

### Establishment of primary cultures

Kidney cell lines were established from kidney tissue obtained from sacrificed animals. Kidney tissue chopped into small (1 mm) pieces and seeded into 6-well plates in DMEM supplemented with 10% FCS and penicillin, streptomycin, gentamycin, nystatin and amphotericin B. After outgrowth in the course of about two weeks, cells were transferred to 25cm^2^ and later 75cm^2^ flasks.

### Virus

The primate herpesviruses CeHV2 (SA8) strain B264 and PaHV2 (HVP2) strain X313 were a kind gift by David Brown and Matthew Jones, Public Health England. ZIKV strain SPH2015 was cloned from RNA and recovered from transfected Vero76 cells (24). Viruses were propagated on Vero76 cells by low multiplicity of infection (MOI) and harvested when complete cytopathic effect had developed.

The recombinant Vesicular stomatitis virus VSV ncp* harboring four amino acid changes associated with reduced cytotoxicity (ncp) and eGFP as reporter (the asterisk stands for eGFP) has previously been described (25).

### Retroviral transduction

For production of Lentivirus-based transducing virus, 293T cells were seeded in T25 flasks (at about 10^6^ cells/flask) and transfected on the next day with 6 μg vector pLenti-IHy-TERT along with 3 μg HIV gag-pol(SCA) and 3 μg VSV-G expression plasmid (pHIT/G), employing the Ca-phosphate co-precipitation method as described before for MLV-based transduction (26, 27). Three days after transfection, cell culture supernatants were harvested, filtered through 0.45 μm filters and stored at −80°C.

For transduction and selection we followed a recently established protocol (28). Thus, cells were seeded in a 96 well plate at 5.000 cells per well. On the next day, 50 μl of transducing virus was added followed by spinoculation at 4,000 × g for 30 min. After two days, cells were transferred into a 24 well plate and selection medium containing 250 μg/mL hygromycin.

### Infection with retroviral pseudoparticles

Retroviral pseudoparticles were produced on 293T cells essentially as described for transducing retroviruses. Cells were transfected with 6 μg vector MLV-luc, 3 μg MLV-gag-pol and 3 μg expression plasmid for the desired viral glycoprotein. Harvested supernatants filtered through 0.45 μm filters and stored at −80°C.

For infection cells were seeded in 96 well plates at 10.000 cells per well. After overnight incubation, medium was replaced with 50 μl fresh medium. For infection 50 μl pseudovirus containing supernatant was added in triplicate samples and incubated for 4 h, after which an additional 100 μl medium were added. After 72 h medium was removed and cells were lysed in 50 μl Luciferase Cell Culture Lysis Reagent (Promega, Madison, WI, USA). Luciferase activity was meastued in a Plate Chameleon V (Hidex, Turku, Finland) microplate reader employing Beetle-Juice (PJK Biotech, Kleinblittersdorf, Germany) as substrate.

### Induction of interferon system

For analysis of the interferon system, cells were seeded in 12-well plates and either treated with interferon (IFN) or infected with VSV ncp*, which induced high amounts of IFN (25). For IFN treatment, cells were incubated for 24 h with medium containing 100 U/mL universal type I IFN-α (pan-Interferon; 11200-2, PBL Assay Science, Piscataway), a chimeric interferon constructed from human IFN alpha A and alpha D (29). For infections cells were inoculated with medium containing VSV ncp* at MOI 0.1. After 24 h cells were harvested for RNA isolation and quantitative PCR.

### Virus replication kinetics and titration

For single step growth kinetics cells were seeded in 24 well plates at 50.000 cells/well and infected on the next day with MOI 1 of the respective viruses. For infection, medium was removed and 500 μl inoculum added for an 1 h incubation at 37°C. After removal of inoculum, cells were washed with phosphate-buffered saline (PBS) and finally incubated in 500 μl culture medium. Cell culture supernatant was harvested at fixed time points (1, 24, 48, 72 and 96 h post infection), centrifuged at 4000 rpm for 5 min to pellet floating cells and the cleared supernatant was frozen at −80°C.

Titration of herpesviruses by plaque assay was essentially performed as described previously (30).

Titers of ZIKV were determined in a modified focus forming assay originally developed for IAV (26, 31). Briefly, Vero76 cells were seeded in 96 well plates at 20.000 cells per well. On the next day, medium was removed and 100μl of tenfold dilutions of virus containing supernatant was added for 1 h at 37°C. After removal of inoculum, 100 μl DMEM medium containing 1% methyl cellulose was added and cells incubated for 72 h at 37°C. Then, the methyl cellulose containing medium was removed and cells washed 1-2 times with PBS. Cells were fixated with cold methanol (10 min −20°C), dried and, after rehydration with PBS, quenched (PBS, 0.5% triton, 20mM glycin) and blocked (PBS, 0.5% triton, 1% BSA). Subsequently, cells were incubated with 50 μl/well hybridoma 4G2 supernatant followed by 50μl secondary anti-mouse horseradish peroxidase (HRP)-conjugated antibody (1:1000, Dianova, Hamburg, Germany). Antibody incubation steps were followed by washing steps. Finally, wells were reacted with TrueBlue peroxidase substrate (Seracare, Milford, MA, USA) until blue foci developed, which were counted, Viral titers were calculated and expressed as focus forming units per milliliter (ffu/mL).

### Quantitative real-time PCR

RNA for quantitative real-time PCR (qRT-PCR) was isolated using the RNeasy Mini Kit (Qiagen, Hilden, Germany) according to the protocol of the manufacturer and eluted in a final volume of 25 μl RNase-free water. To remove contaminating DNA, 1 μg RNA was treated with 0.5 U RNase-free DNase I for 10 min and the reaction stopped by addition of EDTA (final concentration 5 mM) and heating to 75°C for 10 min. For cDNA synthesis 8 μl of DNase-digested RNA were reverse transcribed using random hexamers and the SuperScript™ III First-Strand Synthesis System (Thermo, Waltham, MA, USA) according to the protocol of the manufacturer. Then 1 μl of the cDNA preparation were used for quantitative PCR using a QuantiTect SYBR Green PCR kit (Qiagen, Hilden, Germany) and the Rotorgene Q platform (Qiagen, Hilden, Germany). Primers against MX1 (forward 5’-TTCAGCACCTGATGGCCTATC-3’, reverse 5’-TGGATGATCAAAGGGATGT-GG-3’), IFNB1 (forward 5’-CAGCAATTTTCAGTGTCAGAAGC-3’, reverse 5’-TCATCCTGTCCTTGAGGCAGT-3’) and the housekeeping gene 18S rRNA (forward 5’-GATCCATTGGAGGGCAAGTCT-3’, reverse 5’-CCAAGATCCAACTACGAGCTT-3’) were previously described (32–34). Induction of MX1 and IFNB1 expression was analyzed by the 2^−ΔΔCT^ method (35) using 18S rRNA as reference gene.

### Flow cytometry

For flow cytometry (FCS) 10^6^ cells/well were seeded in 6-well plates, grown overnight and detached with accutase (Invitrogen, Carlsbad, CA, USA). After centrifugation (1200 rpm, 5 min) cells were resuspended and washed once in FCS buffer (PBS, 5 % FCS, 2 mM EDTA). After removing the supernatant cells were stained with a rat anti-podoplanin antibody (1:100; Origene AM01133PU-N, Rockville, MD, USA) or no antibody in a volume of 50 μl for 30 min on ice. After three washing steps with FCS buffer, cells were reacted with secondary Alexafluor488-conjugated donkey anti-rat antibody (1:100; Invitrogen, Carlsbad, CA, USA). After three washes in FCS buffer cells were fixated in 2% PFA. Cells were analyzed in a LSR II flow cytometer (Becton, Dickinson, East Rutherford, NJ, USA) using FACS Diva software. Diagrams for figures were prepared using FCS Express 4 (De Novo Software, Pasadena, CA, USA).

### Microscopy

Brightfield images were taken at 10x magnification (10x/0.45 Plan-Apochromat objective) on a Zeiss Axio Observer Z1/7 equipped with an Axicom 503 using ZEN software. Images were cropped and further processed (adjustment of brightness, scale bar) using ImageJ/Fiji (36).

## Results

### Establishment of immortalized rhesus macaque kidney cell lines

To establish cell lines from rhesus macaques we dissociated kidney tissues from adult male (6 years) and female (9 years) animals and established primary cultures of kidney cells. The cell cultures were named *Macaca mulatta* kidney (MaMuK) cell lines MuMuK2345C, MaMuK2345MW, and MaMuK8639. After several passages cells were transduced with the human TERT gene, which has been shown to immortalize cells. For efficient transduction an HIV-1-based lentiviral system was modified to include a chimeric Gag protein to avoid TRIM5α-mediated restriction (21, 22). By selection with puromycin, resistant cell cultures were established. Compared to the parental cultures the transduced cultures showed faster growth and a smaller cell body (Fig. 1). All cell lines displayed a spindle-like morphology as seen in mesenchymal cells. Cells were successfully passaged at least 30 times, during which the cell population doubled at least 70 times. Expression of TERT in transduced but not parental cells was confirmed by Western blot (Fig. 2). Finally, flow cytometric analysis of TERT-positive cells revealed expression of podoplanin (PDPN) (Fig. 3), a podocyte maker (37) that is also expressed on 293T cells (38), a human embryonic kidney cell line frequently used for biomedical research. Thus, we had obtained three TERT and PDPN expressing permanent rhesus macaque kidney-derived cell lines that were subjected to further analysis.

**Figure 1:**
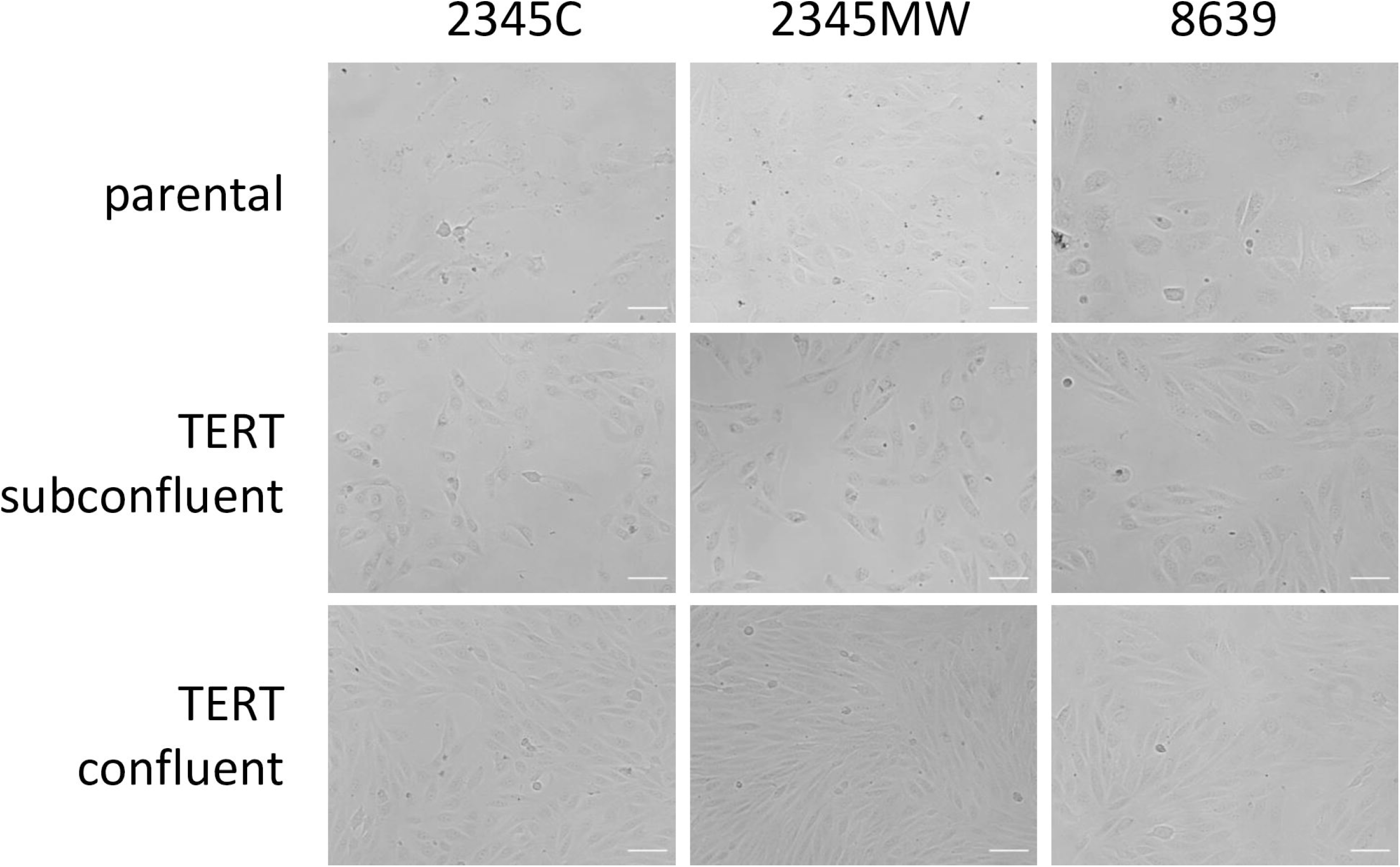
Morphology of parental and TERT immortalized rhesus kidney cells. Cells were seeded in 6-well plates and bright field images taken at 10x magnification. White scale bars indicate 100 μm.

**Figure 2:**
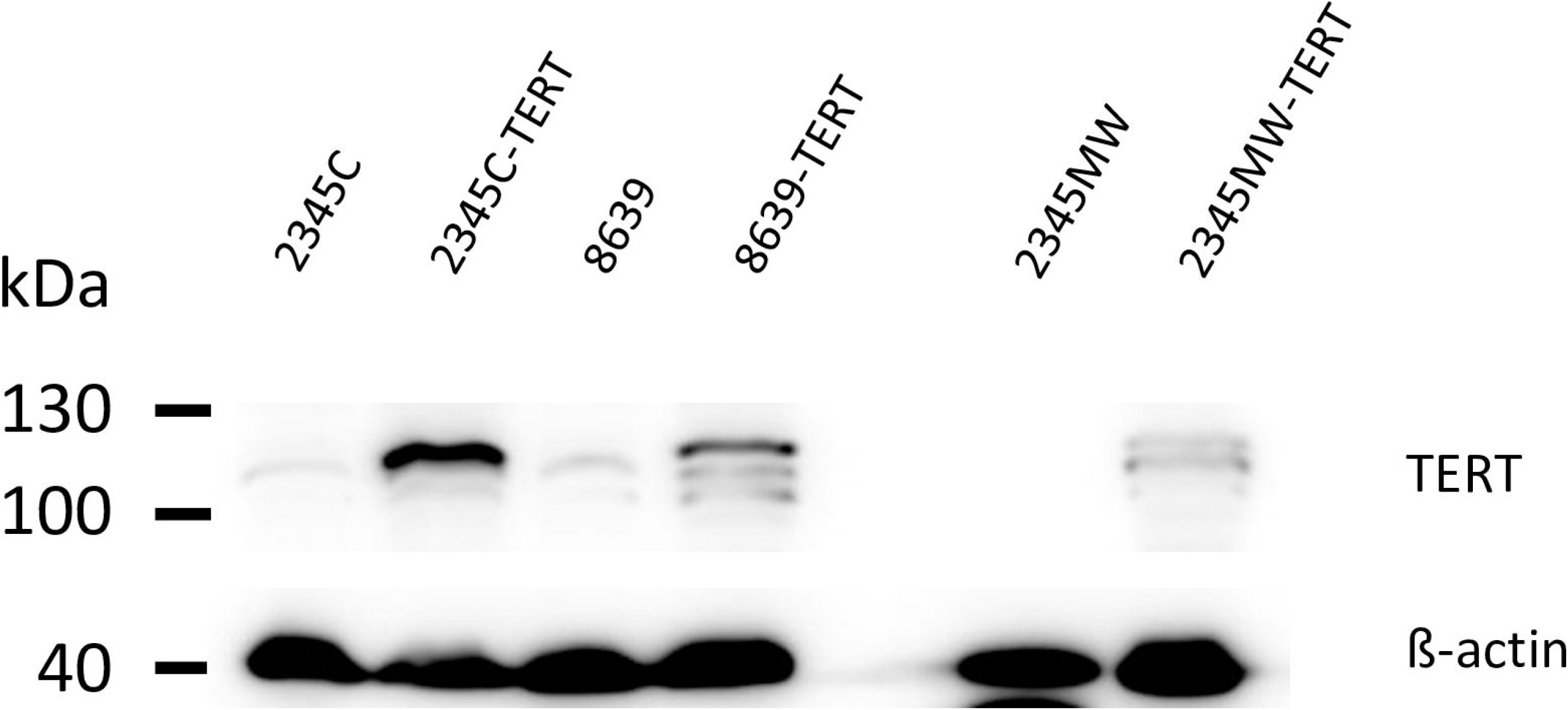
Immortalized kidney cells express transduced immortalization gene TERT. Lysates of parental and immortalized kidney cells were analyzed in western blot for expression of immortalization gene TERT. Detection of β-actin (ACTB) served as loading control. The results were confirmed in an independent experiment.

**Figure 3:**
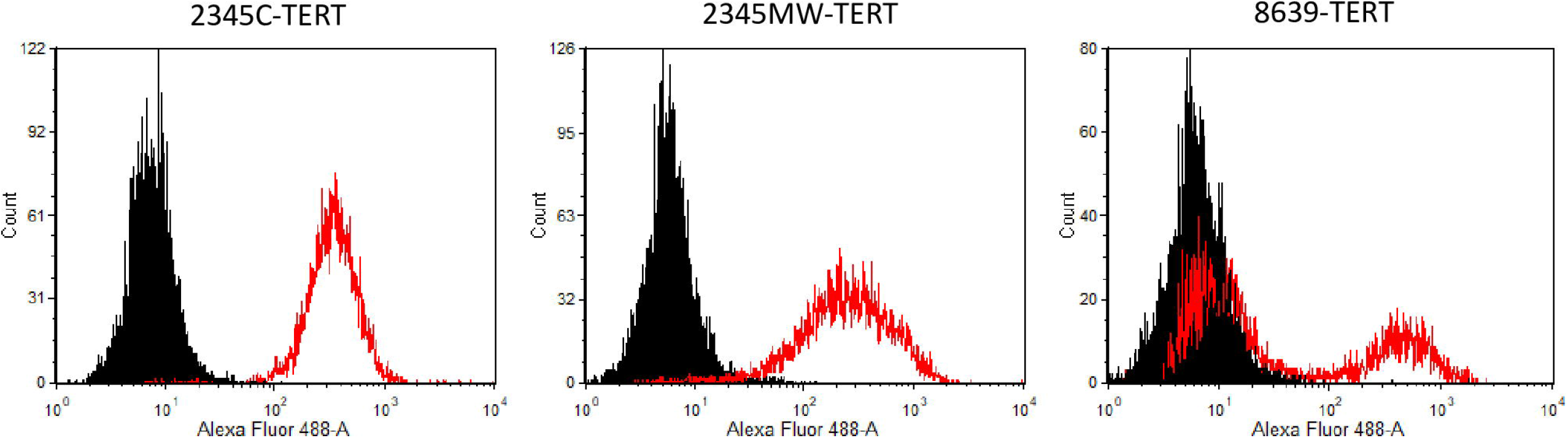
Immortalized kidney cells express the surface marker podoplanin. Cells were detached and stained with anti-podoplanin antibody or no antibody and subjected to flow cytometry.

### Immortalized rhesus macaque kidney cell lines have a functional IFN system

We next characterized whether the cells expressed IFN-stimulated genes (ISGs) in response to IFN or virus infection. For this, we treated the cells either for 24 h with 100 U/mL pan-IFN, a chimeric human IFN alpha, or vesicular stomatitis virus (VSV) ncp*, which strongly induces IFN (25). Induction of interferon β (IFNB1) or MX1 was analyzed by quantitative RT-PCR. The human lung cell line A549 was included as positive control, since this cell line has an intact IFN system (39). Infection with VSV but not treatment with pan-IFN strongly induced IFNB1 expression in all rhesus macaque kidney cell lines analyzed and induction efficiency was roughly comparable to that measured for A549 cells (Figure 4a). Both, infection with virus and treatment with pan-IFN, strongly induced MX1 expression in the rhesus macaque kidney cell lines (Figure 4b). Levels of MX1 induction were lower than in A549 cells for all MaMuK cells but still more than 300-fold over background (Figure 4b). Collectively, these results indicate that all newly established cell lines had an intact IFN system.

**Figure 4:**
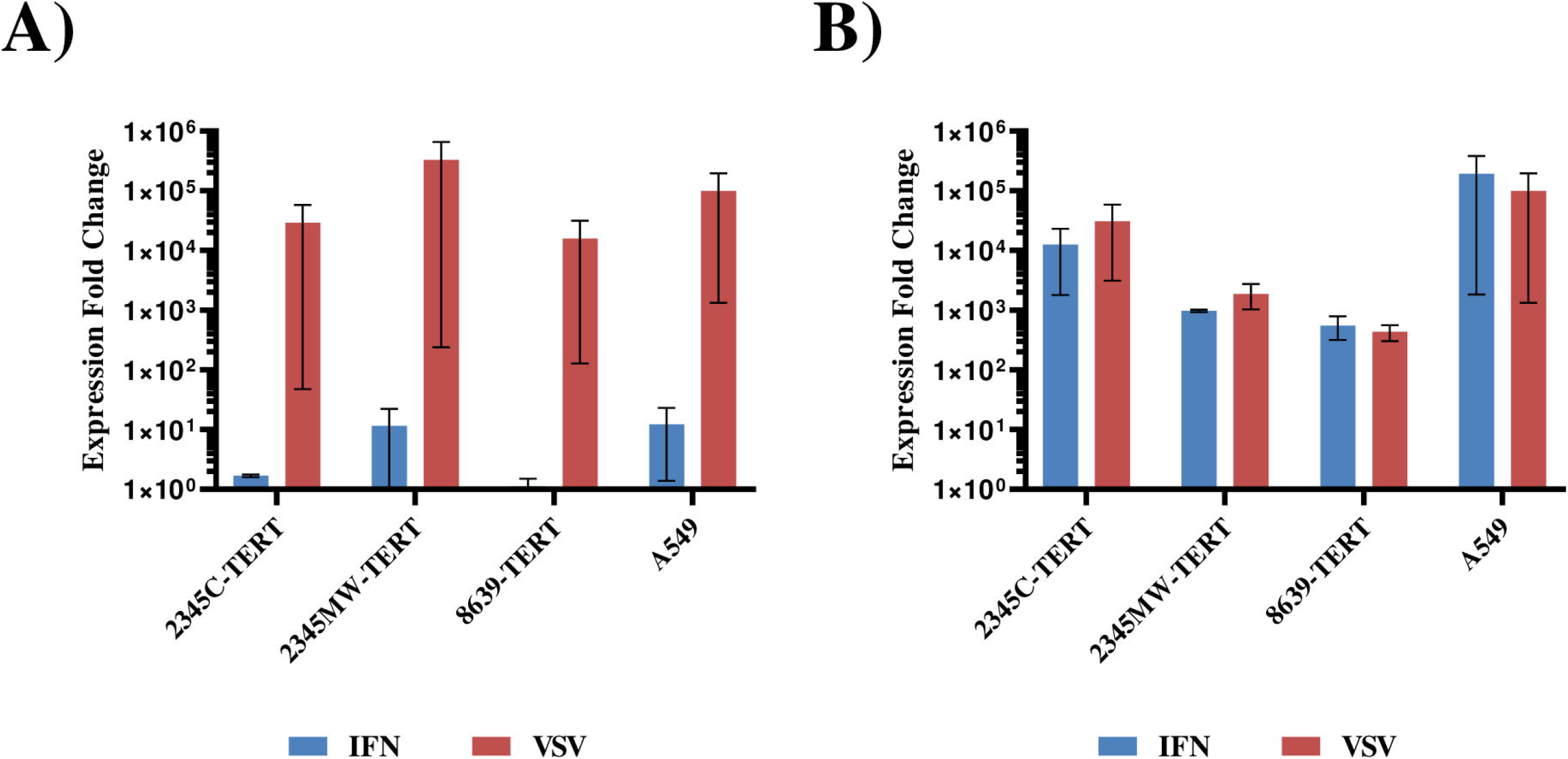
Immortalized kidney cells have a functional interferon system. Cells were seeded in 12 well plates and either treated with 100 U/mL pan-interferon or infected with VSV ncp* (MOI 0.1). Untreated cells served as control. After 24 h cells were harvested, RNA was isolated and analyzed by quantitative RT-PCR for expression of (A) interferon beta (IFNB1) or (B) MX1. Transcript levels were normalized against 18S rRNA transcript levels and expression fold change calculated with respect to control cells. The average of two independent experiments is shown. Error bars indicate standard error of the mean.

### Immortalized rhesus monkey kidney cells lines are susceptible and permissive for virus infection

We next analyzed if these cell lines were susceptible and permissive to viral infection. First, we chose a retroviral pseudotyping system to study susceptibility of cells to entry driven by the glycoproteins from Indiana vesiculovirus (VSV), Ebola virus (EBOV), Nipah virus (NIV), influenza A virus (IAV) or Lassa virus (LASV). Viral particles bearing no glycoprotein served as negative control while 293T target cells served as positive control since this cell line is known to allow entry driven by the above listed glycoproteins. The rhesus macaque kidney cell lines allowed for entry driven by all glycoproteins tested and entry efficiency was comparable to that measured for 293T cells (Figure 5). In some cases, the TERT-immortalized cell lines demonstrated higher luciferase activity (Figure 5) relative to 293T cells, which may be related to their higher growth rate.

**Figure 5:**
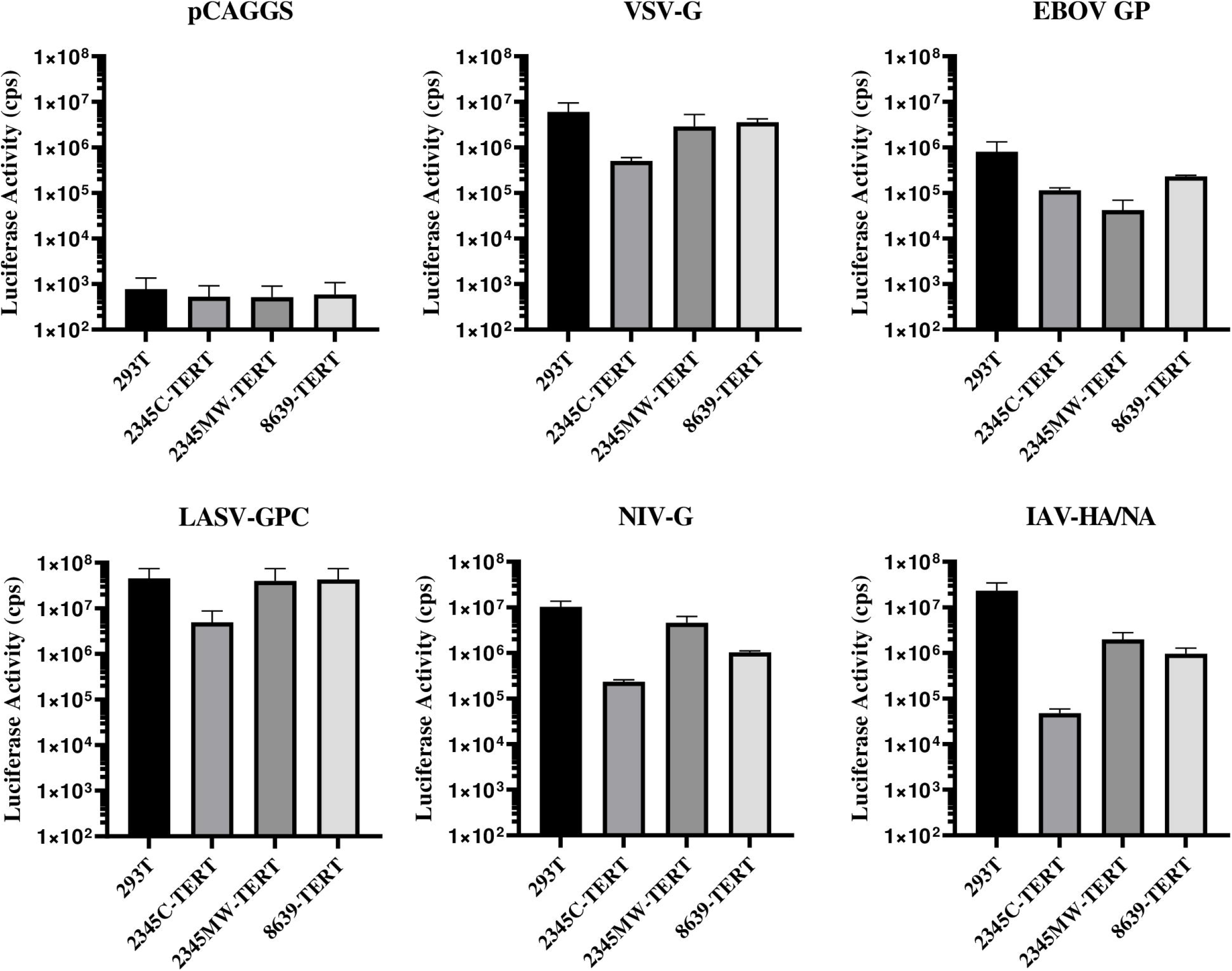
Immortalized rhesus kidney cells are susceptible to entry driven by diverse viral glycoproteins. Parental and immortalized rhesus kidney cells were seeded in 96 well plates and transduced in triplicates with MLV pseudoparticles encoding firefly luciferase and bearing the indicated viral glycoproteins. Transduction of 293T cells served as positive control while pseudoparticles without viral glycoprotein served as negative control. After 72 h cell lysates were harvested and luciferase activities determined. The average of two independent experiments is shown. Error bars indicate standard error of the mean.

Finally, we determined whether the rhesus macaque kidney cell lines were permissive to infection with selected RNA and DNA viruses. As DNA viruses we employed the primate simplexviruses Papiine alphaherpesvirus 2 (PaHV2) and Cercopithecine alphaherpesvirus 2 (CeHV2), which naturally infect NHP, and as RNA virus we used ZIKV, a human pathogen. ZIKV replicated in all macaque cell lines with similar efficiency as compared to Vero cells (Figure 6c), which are routinely used to amplify ZIKV. The two herpesviruses showed a differential behavior. While CeHV2 replicated with markedly reduced efficiency in the rhesus macaque kidney cell lines as compared to Vero76 cells (Figure 6b), PaHV2 replicated to high levels in the macaque cell lines (Figure 6a).

**Figure 6:**
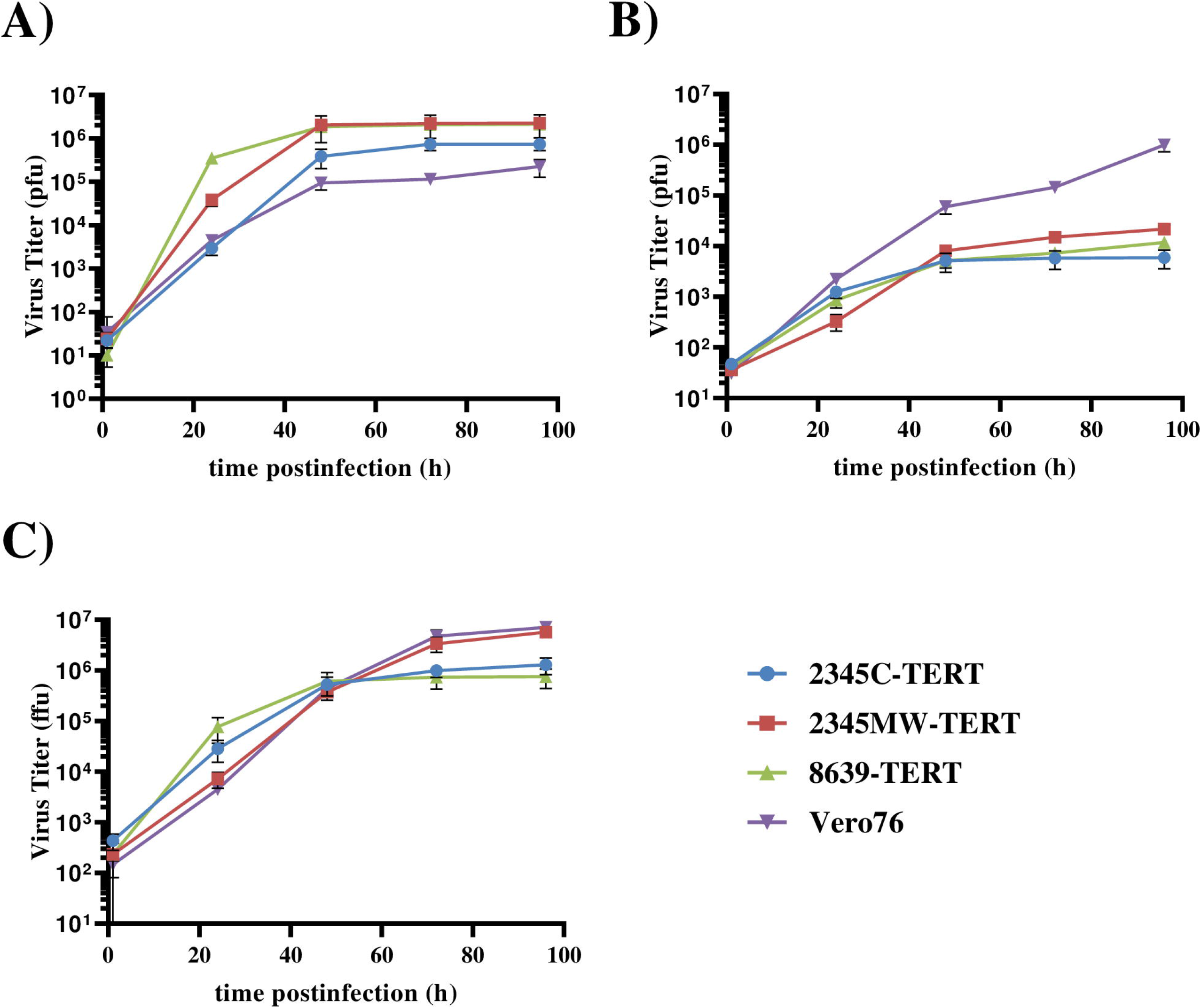
Immortalized rhesus kidney cells support growth of primate herpesviruses and Zika virus. Cells were seeded in 24 well plates and triplicate samples were infected with (A) Papiine alphaherpesvirus 2, (B) Cercopithecine alphaherpesvirus 2 or (C) ZIKV strain SPH2015 at an MOI of 1. Vero76 cells served as positive control. Supernatants were harvested after 1, 24, 48, 72 and 96 h postinfection and infectious virus titer was determined by plaque assay (herpesviruses) or focus formation assay (ZIKV) on Vero76 cells. Virus titers are shown as plaque forming units (pfu) or focus forming units (ffu), respectively. The results of a single representative experiment are shown and were confirmed in an independent experiment. Error bars indicate standard deviation.

## Discussion

We generated and characterized three cell lines derived from rhesus macaque kidney, which were immortalized by human TERT. These cells were susceptible to infection by diverse viruses, as determined with pseudotyped particles, and were permissive to infection with primate simplexviruses and ZIKV. Currently, a single rhesus macaque kidney cell line, LLC-MK2, is available from cell repositories. LLC-MK2 cells have epithelial morphology, whereas our cell lines display a spindle-like morphology as seen in mesenchymal cells. By flow cytometry, we could detect expression of the marker podoplanin (PDPN), placing them into the podocyte lineage according to single cell RNAseq data on kidney (37).

For analysis of viral infections, the status of the IFN system is of importance since sensing of viruses and subsequent induction of ISG expression can efficiently limit viral replication. As a consequence, cell lines that allow efficient amplification of diverse viruses, like Vero cells, frequently harbor a defective IFN system (40, 41). We could demonstrate that the new cell lines have an intact IFN system as evidenced by strong IFN beta induction upon VSV infection and strong induction of the ISG MX1 by IFN or viral infection.

The rhesus macaque cells lines allowed entry driven by the glycoproteins from representatives of several virus families such as Rhabdoviridae, Filoviridae, Arenaviridae and Paramyxoviridae. Entry driven by the IAV (Orthomyxoviridae) hemagglutinin was also detected, but was less efficient than for human 293T cells. As a consequence, we expect that the cell lines should be susceptible to infection by diverse viruses. Indeed, productive infection could be demonstrated for two primate herpesviruses, PaHV2 and CeHV2, as well as ZIKV (Flaviviridae). Virus titers were comparable or even higher than those attained in a highly permissive control cell line, Vero, despite a functional IFN system. Growth of CeHV2 in the rhesus macaque kidney cells was less efficient than in Vero cells, which may be due to species specific restrictions, as observed earlier (30).

## Conclusions

In summary, we established three TERT-immortalized rhesus macaque kidney cell lines with an intact IFN system, which might support infection by diverse viruses, including primate herpesviruses and Zika virus. These cell lines could be valuable tools in comparative infection research or in translational research.

## Acknowledgments

We would like to thank Christiane Stahl-Hennig, Alexander von Hahn (DPZ, Germany), Andrea Maisner (Philipps University of Marburg, Germany), Michael Farzan (The Scripps Research Institute, USA), Parmjit Jat (Ludwig Institute for Cancer Research, UK), David Brown and Matthew Jones (Public Health England, UK) for their kind gift of material.

